# Single-cell analysis of CX3CR1^+^ cells reveal a pathogenic role for BIRC5^+^ myeloid proliferating cells driven by *Staphylococcus aureus* leukotoxins

**DOI:** 10.1101/2023.02.27.529760

**Authors:** Denis G. Loredan, Joseph C. Devlin, Keenan A. Lacey, Nina Howard, Ze Chen, Erin E. Zwack, Jian-Da Lin, Kelly V. Ruggles, Kamal M. Khanna, Victor J. Torres, P’ng Loke

## Abstract

Our previous studies identified a population of stem cell-like proliferating myeloid cells within inflamed tissues that could serve as a reservoir for tissue macrophages to adopt different activation states depending on the microenvironment. By lineage tracing cells derived from CX3CR1^+^ precursors in mice during infection and profiling by scRNA-seq, here we identify a cluster of BIRC5^+^ myeloid cells that expanded in the liver during either chronic infection with the parasite *Schistosoma mansoni* or the bacterial pathogen *Staphylococcus aureus*. In the absence of tissue damaging toxins, *S. aureus* infection does not elicit these BIRC5^+^ cells. Moreover, deletion of BIRC5 from CX3CR1 expressing cells results in improved survival during *S. aureus* infection. Hence, the combination of scRNA-Seq and genetic fate mapping CX3CR1^+^ cells revealed a toxin dependent pathogenic role for BIRC5 in myeloid cells during *S. aureus* infection.

## Introduction

During inflammation, CX3CR1^+^ monocytes are recruited into inflamed sites and can respond to signals to adopt different activation states when exposed to different environmental stimuli in the tissues (1,2). We had previously used a combination of single-cell RNA sequencing (scRNA-seq) and genetic fate mapping to profile cells derived from CX3CR1^+^ precursors in mice during atherosclerosis (3). The results revealed a spectrum of macrophage activation states more complex than the traditional M1 and M2 categories in the inflamed plaques (3). Additionally, we identified an unexpected cluster of proliferating myeloid cells with a stem cell–like signature that we hypothesize may persist in a proliferating self-renewal state in inflamed tissue instead of differentiating immediately into macrophages after entering the tissue. We speculated that these proliferative cells may serve as a reservoir for tissue macrophages that adopt different activation states during an inflammatory response. However, it was unclear if this population was unique to atherosclerosis or could be observed in other inflammatory conditions such as during parasite and bacterial infections.

During infection with the helminth parasite *Schistosoma mansoni*, macrophages derived from Ly6C^high^ CX3CR1^+^ monocytes are required to protect the liver hepatocytes from tissue damage as part of the granulomas that are formed around the parasite eggs (4,5). This chronic inflammatory response in the liver has been well characterized as a model of monocyte recruitment and differentiation into activated macrophages important for the prevention of tissue damage (6). *Staphylococcus aureus* is a Gram-positive bacterium that produces a collection of pore-forming leukocidins as a strategy for immune evasion and tissue damage (7,8). Leukocidins are important virulence factors that target and kill a variety of immune cells and *S. aureus* mutants that lack leukocidins are attenuated (9–12). However, the exact mechanisms by which leukocidins promote pathogenesis during *S. aureus* infection is still incompletely understood and there is a possibility that increased inflammatory responses driven by these toxins as immune subversion molecules may be damaging instead of being protective to the host. Here, we investigated if the proliferative myeloid cells that we first observed in the context of atherosclerosis are also present in the liver during *S. mansoni* and *S. aureus* infections.

BIRC5, also termed “Survivin”, is an important protein for cell division and inhibition of apoptosis (13,14). It is most well characterized in the context of cancer cells, where it has received considerable attention as a potential drug target (15–17). This small adaptor protein interacts with its cellular partners during mitosis as part of the chromosomal passenger complex (CPC) as well as with members of the inhibitors of apoptosis (IAP) protein family to interfere with apoptosis (14). The BIRC5 inhibitor YM155 has been shown to have anti-cancer effects in cell culture and in mouse tumor models, but has yet to be utilized as an effective clinical drug (17,40–41). A potential role in autoimmune diseases such as rheumatoid arthritis (18) and multiple sclerosis (19,20) as well as in CD4^+^ cells during HIV-1 infection (21) has also been described.

While the role of BIRC5 has been well characterized at a cellular level in cancer cells, the role played by BIRC5 in myeloid cells during infection is not well understood. By utilizing singlecell RNA-Seq with genetic fate mapping to characterize immune cells derived from CX3CR1^+^ precursors in the liver during *S. mansoni* and *S. aureus* infection, we identified and further investigated a population of BIRC5^+^ myeloid proliferating cells with stem cell–like signatures, which may play a pathogenic role during infection with *S. aureus*. Hence, small molecules designed to inhibit the activity of BIRC5 may be beneficial as host-direct therapy for *S. aureus* infection.

## Materials and Methods

### Mice

B6.*Cx3cr1*^CreERT2-IRES-EYFP/+^ were generously provided by D. Littman (Department of Pathology, Department of Microbiology, NYU Langone Health, New York, NY 10016). *B6.Rosa26*^stop-tdTomato^ mice (JAX: 007914) were from Jackson Laboratories (Bar Harbor, ME).

B6.*Cx3cr1*^CreERT2-IRES-EYFP/+^ mice and B6.*Rosa26*^stop-tdTomato^ mice were crossed to generate *Cx3cr1*^CreERT2-IRES-EYFP/+^ *Rosa26*^tdTomato/+^ mice as previously described. B6.129P2-*Birc5^tm1Mak^*/J were purchased from Jackson Laboratories (strain #031830), and crossed with B6.*Cx3cr1*^CreERT2-IRES-EYFP/+^ to generate *Cx3cr1*^CreERT2-IRES-EYFP/+^ *Birc5*^Flox/Flox^ conditional knockout mice (37). To induce labeling, Tamoxifen (Sigma-Aldrich) was dissolved in corn oil and given by oral gavage at a dose of 500 mg/kg body weight. Alternatively, mice were placed on Tamoxifen diet (Teklad Global, 250, 2016, Red) with diet code TD130856, to induce conditional deletion. All animal procedures were approved by the NYU School of Medicine IACUC Committee.

### Models of Infection

For experiments with *Schistosoma mansoni* infection, mice were infected percutaneously with 75 cercariae, given an oral gavage of Tamoxifen at 6.5-7 weeks post-infection, and analyzed at 8 weeks post-infection. The reagent was provided by the NIAID Schistosomiasis Resource Center of the Biomedical Research Institute (Rockville, MD) through NIH-NIAID Contract HHSN272201700014I. NIH: *Biomphalaria glabrata* (NMRI) exposed to *Schistosoma mansoni* (NMRI).

*Staphylococcus aureus* strains USA300 LAC clone AH1263 (wildtype) (38) and AH-LAC Δ*lukAB, hlg::tet, lukED::kan,pvl::epc, hla::erm* (ΔTOX) were used in this study (42). For *in vivo* infection studies, wildtype *S. aureus* was grown on tryptic soy agar (TSA) or tryptic soy broth (TSB). *S. aureus* was grown overnight and a 1:100 dilution of overnight cultures was sub-cultured into fresh TSB. *S. aureus* grown to early stationary phase (3 h) was collected and normalized by OD600 for further experimental analysis. Mice were anesthetized with Avertin (2,2,2-tribromoethanol dissolved in tert-Amyl alcohol and diluted to a final concentration of 2.5 % vol/vol in 0.9 % sterile saline) by intraperitoneal injection. For the bloodstream infection model, mice were challenged with 1×10^7^ CFU by retro-orbital injection, after being placed on tamoxifen diet 2 days prior. Survival was assessed by continuous monitoring and a 30% weight loss cutoff for humane euthanasia. To evaluate bacterial burden in organs, infected mice were sacrificed, and the indicated organs collected in 1 ml PBS and homogenized to enumerate bacterial burden. Blood was collected by cardiac puncture and anti-coagulated using heparin coated tubes.

For sorting and single-cell RNA-sequencing, mice were infected by retro-orbital injection with either 5×10^6^ CFU wildtype or ΔTOX *S. aureus*, and given an oral gavage of Tamoxifen just prior to infection. Infected mice were analyzed at 7 days post-infection.

### Liver Immune Cell Isolation

Liver tissues were processed as described. Livers were chopped and incubated in RPMI + 10% FBS with collagenase VIII (100U/mL; Sigma Aldrich) and DNase I (150 μg/mL; Sigma Aldrich) for 45 minutes at 37° C and then passed through a 70-μm cell strainer (Fisher Scientific). Leukocytes were enriched for by density-gradient centrifugation over a 40/80% Percoll (GE Healthcare) gradient, and remaining red blood cells were lysed with ACK lysis buffer (Lonza) and washed in PBS and used for cell sorting or flow cytometry analysis.

### Flow Cytometry and Cell Sorting

Single-cell suspensions were stained with fluorescently conjugated antibodies in a 1:100 dilution unless otherwise noted. Cells were stained with LIVE/DEAD Blue Reactive Dye (Invitrogen) and 4μg/mL anti-CD16/32 (2.4G2; Bioxcell). The following anti-mouse antibodies were used, with clone and source company listed: CD45 PerCP-Cyanine5.5 (30-F11, BioLegend), CD45 BV510 (30-F11, BioLegend), CX3CR1 BV605 (SA011F11, BioLegend), F4/80 BV711 (BM8, BioLegend), Ly6G PerCP-Cyanine5.5 (1A8, BioLegend), Ly6C PE/Cyanine7 (HK1.4, BioLegend), CD11b BUV395 (M1/70, BD Horizon), CD11c BV650 (N418, BioLegend), Ki67 PE/Dazzle (16A8, BioLegend). Intracellular Ki67 was stained using the eBioscience Transcription Factor Staining Kit, following the included protocol. The BD FACS ARIA II was used for cell sorting, and the Bio-Rad ZE5 Yeti was used for flow cytometry analysis. Data analysis was performed using FlowJo v10 (FlowJo LLC).

### Single-cell RNA-sequencing

Single cell suspensions were obtained from livers as described above. For each data set, cells isolated from at least 3 mice were pooled together, with equal numbers of cells pooled from each animal used. 12,000 cells from each group were loaded on a 10X Genomics Next GEM chip and single-cell GEMs were generated on a 10X Chromium Controller. Subsequent steps to generate cDNA and sequencing libraries were performed following 10X Genomics’ protocol. Libraries were pooled and sequenced using Illumina NovaSeq SP 100 cycle as per 10X sequencing recommendations.

The sequenced data were processed using Cell Ranger (version 6.0) to demultiplex the libraries. The reads were aligned to *Mus musculus* mm10 and SCV2 (MN985325.1) genomes to generate count tables that were further analyzed using Seurat (version 4.1.2). Data are displayed as uniform manifold approximation and projection (UMAP).

### Cytokine Analysis

Cytokine quantification was determined using the MILLIPLEX MAP Mouse Cytokine/Chemokine Magnetic Kit (MCYTMAG-70K-PX32, Millipore). The assay was conducted as per the manufacturer’s instructions and plates were ran on a MAGPIX instrument with xPONENT software. Statistical analyses were performed in Prism for each individual cytokine.

## Results

### Single-cell RNA-Seq analysis of immune cells derived from CX3CR1^+^ precursors during *Schistosoma mansoni* infection

During infection with *S. mansoni*, extracellular parasite eggs lodged in the liver induce the recruitment of CX3CR1^+^ monocytes to form a granuloma as part of the host response in limiting tissue damage and hepatotoxicity (4,6). To perform scRNA-seq on immune cells derived from CX3CR1^+^ precursors, we made use of a previously described fate-mapping mouse model in which *Cx3cr1*^CreERT2-IRES-EYFP^ mice were crossed with *Rosa26*^stop-tdTomato^ reporter mice (hereinafter referred to as *Cx3cr1*^CreERT2-EYFP/+^ *R26*^tdTomato/+^ mice) (5). Tamoxifen treatment of these mice results in the irreversible labeling of all CX3CR1^+^ cells with tdTomato. We infected *Cx3cr1*^CreERT2-EYFP/+^ *R26*^tdTomato/+^ mice with 75 cercariae of *S. mansoni*, and 17 days prior to analysis at 8 weeks post-infection, mice were administered tamoxifen via oral gavage (Figure 1A). At 8 weeks post-infection, we FACS purified tdTomato^+^ CD45^+^ cells from the livers of infected mice, combined all samples, and submitted 12,000 cells for single-cell RNA-sequencing. Sequencing was performed on the 10X Genomics platform, and cells were visualized by Uniform Manifold Approximation and Projection (UMAP), a machine learning algorithm for dimensionality reduction (Figure 1B). UMAP clustering revealed CX3CR1^+^-derived cells to be a heterogeneous population consisting of both a major myeloid and lymphoid cluster. The myeloid cluster consisted mostly of macrophages characterized by expression of Chil3, Lyz2, and Alox5 while the lymphoid cluster consisted of multiple cell types including CD8^+^ T cells, NK cells, T_H_1, and T_H_2 CD4+ T cells (Figure 1B,C,D). Interestingly, our analysis showed a separate cluster of cells defined by the gene Birc5 that consisted of both myeloid and lymphoid cells (Figure 1B,C,D). This cluster was further defined by the expression of cell cycle and proliferation genes Stmn1 and Ki67 (Figure 1E), similar to the BIRC5^+^ population of monocytes we identified in a previous study (3). These data suggest that a population of BIRC5^+^ stem-cell-like proliferating cluster of myeloid cells is present in the liver during chronic infection with *S. mansoni* and could be a general feature of inflammation, not unique to atherosclerosis (3). However, there are also BIRC5^+^ lymphocytes, which may be derived from a small subset of T cells that can also express CX3CR1 (22,23).

**Figure 1.**
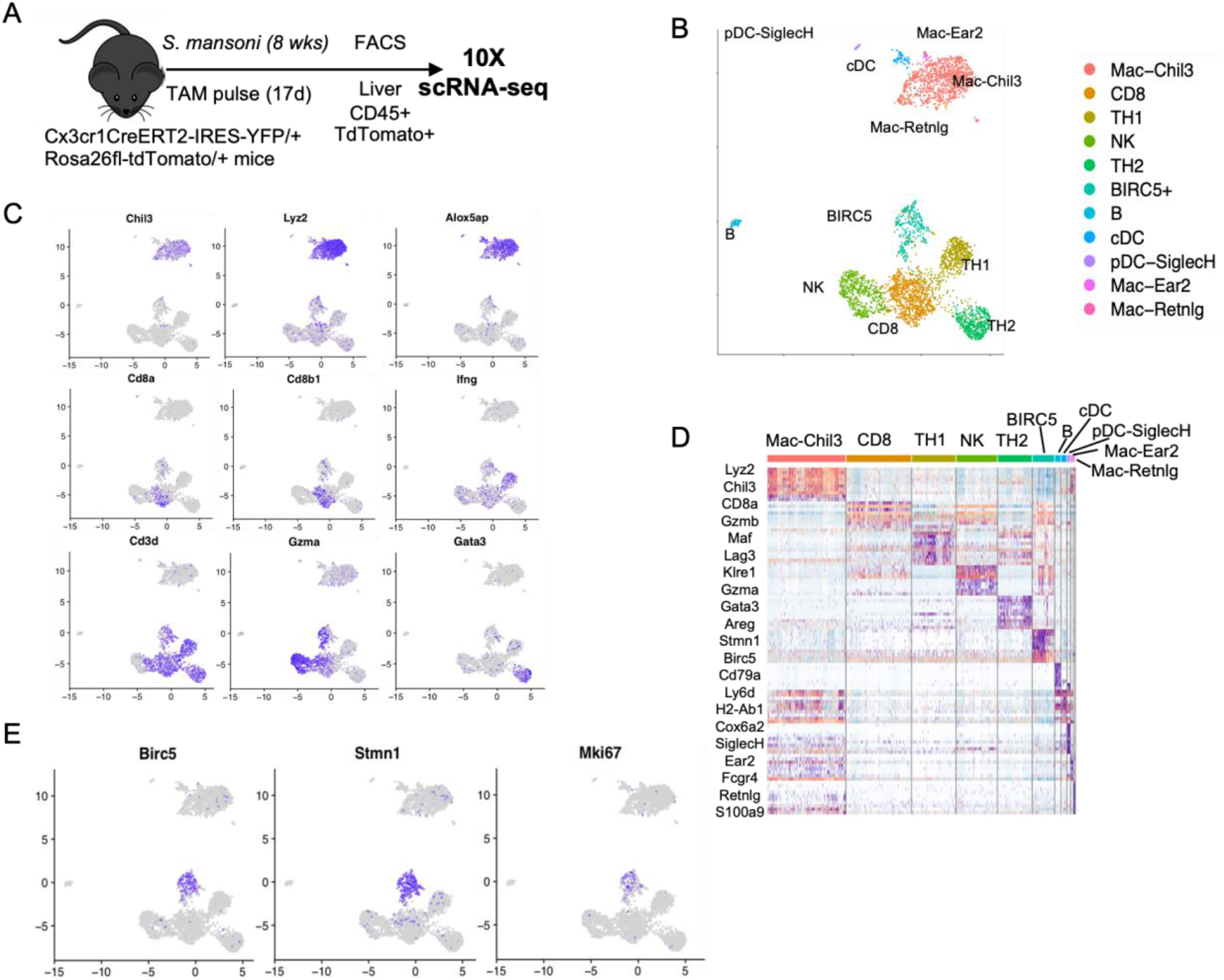
Single-cell RNA-seq of fate-mapped CD45^+^ cells derived from CX3CR1^+^ precursors during *Schistosoma mansoni* infection reveals BIRC5^+^ T cells and BIRC5^+^ myeloid cells. **(A)** Experimental design for single-cell RNA-seq of CD45^+^ from *Cx3cr1*^CreERT2-EYFP/+^ *R26*^tdTomato/+^ mice infected cutaneously with 75 *S. mansoni* and sorted 8 weeks post-infection. Gating was on singlets, live cells, CD45^+^, tdTomato^+^. **(B)** UMAP representation and unbiased clustering of major immune cell populations from the CD45^+^ tdTomato^+^ cells sequenced, with each dot representing individual cells. **(C)** Expression of canonical cellular markers for the major clusters of immune cells. **(D)** Heatmap showing the markers of the main populations of immune cell clusters **(E)** Identification of BIRC5^+^ proliferative cluster.

### ScRNA-Seq analysis reveals leukotoxin dependent BIRC5+ cells during *S. aureus* infection in the liver

We next investigated if the population of BIRC5^+^ myeloid cells derived from CX3CR1+ precursors are also observed in an unrelated infection model characterized by a type 1 immune response in the liver. For this we infected mice with the Gram-positive bacteria *S. aureus*, which is known to infect the liver (26) and would allow us to compare immune cell infiltrates in the same tissue affected by *S. mansoni* infection.

*S. aureus* has evolved to produce virulence factors including leukocidins that target leukocytes and are required for infection (8,12,25). We hypothesized that leukocidins may affect CX3CR1^+^ immune cells during *S. aureus* infection, thus we infected *Cx3cr1*^CreERT2-EYFP/+^ *R26*^tdTomato/+^ mice via retro-orbital injection with 5×10^6^ CFU of either wildtype (WT) *S. aureus*, or a strain in which the genes *hlgABC, lukSF-PVL, lukAB, lukED, and hla* have been deleted, referred to as ΔTOX (Figure 2A). All mice were given an oral gavage of tamoxifen at the time of infection to label cells that express CX3CR1 at that time. At 7 days post-infection we FACS purified tdTomato^+^ CD45^+^ cells from the livers of both WT and ΔTOX infected mice and submitted 12,000 cells each from both groups, separately, for single-cell RNA-sequencing. All cells were visualized together in two-dimensional space using UMAP, with distinct sequencing libraries allowing us to distinguish between cells derived from either WT- or ΔTOX-infected mice (Figure 2B,C). Similar to our results from *S. mansoni*-infected mice, we identified distinct myeloid and lymphoid clusters derived from CX3CR1^+^ precursors in the livers of *S. aureus* infected mice. The myeloid cluster consisted mostly of monocyte and macrophage cell types, however uniquely in this dataset, a neutrophil cluster was also identified. The lymphoid cluster consisted of 3 main cell types: CD8 T cells, NK cells, and NKT cells. Notably, CD4 T cells were absent from this cluster, suggesting the expression of CX3CR1 on CD4 T cells depends on the type of infection, and can occur during helminth infection but not *S. aureus* infection.

**Figure 2.**
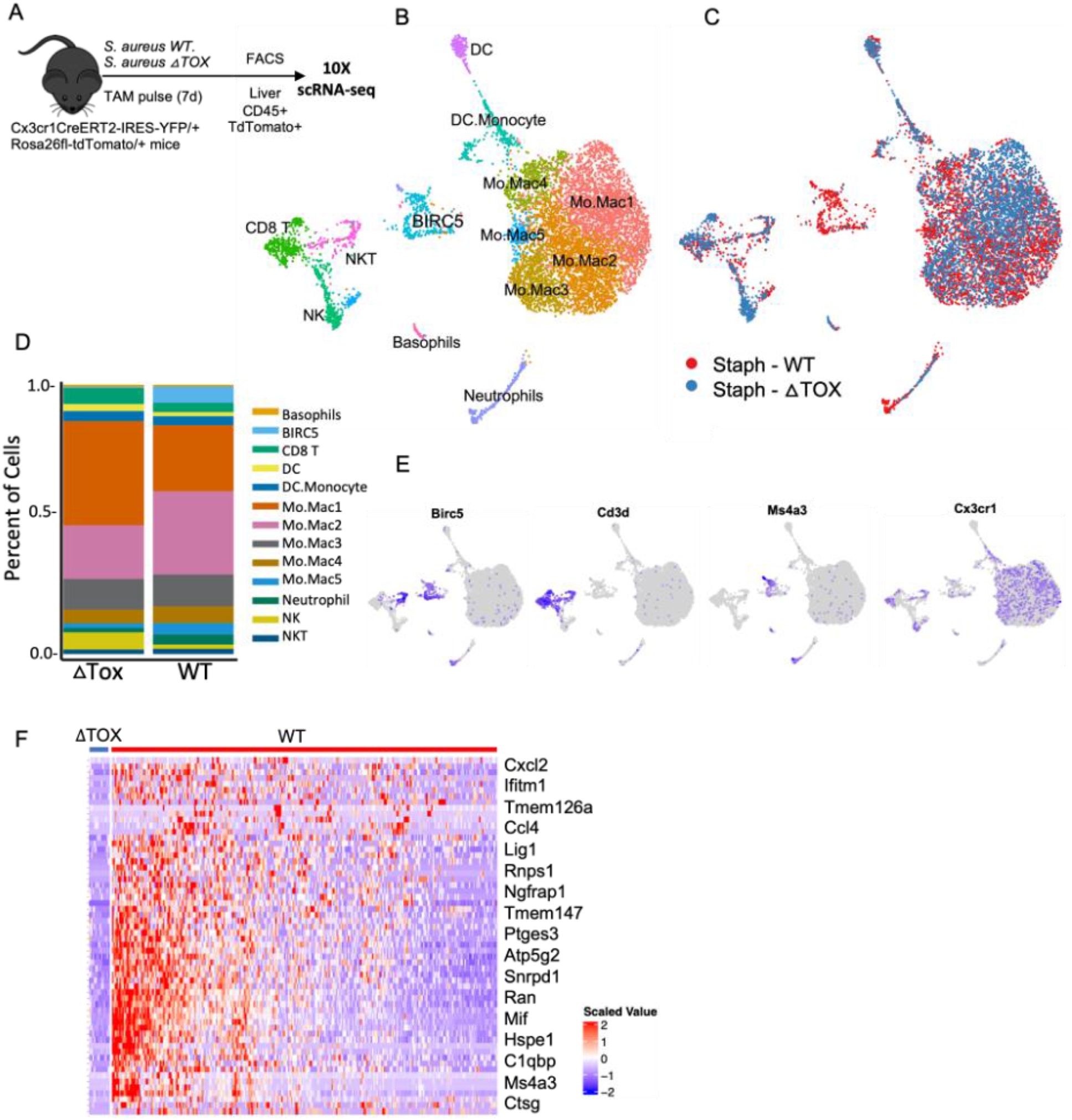
Single-cell RNA-seq reveals BIRC5^+^ immune cells to be specifically elicited by *Staphylococcus aureus* toxins during infection. **(A)** Experimental design for single-cell RNA-seq of CD45^+^ from *Cx3cr1*^CreERT2-EYFP/+^ *R26*^tdTomato/+^ mice infected intravascularly with 5×10^6^ CFU of *S. aureus* and sorted 1 week post-infection. Gating was on singlets, live cells, CD45^+^, tdTomato^+^. **(B)** UMAP clustering of CX3CR1-derived CD45^+^ cells with phenotypic classification**(C)** UMAP clustering of isolated cells differentiating between cells from WT versus ΔTOX infected mice. **(D)** Percentage of total cells defined as each immune subset from either WT or ΔTOX infected mice. **(E)** Expression of BIRC5 and other lineage defining markers across both datasets. **(F)** Heatmap showing differential gene expression between identified immune clusters.

Consistent with our data from *S. mansoni* infection, we saw a distinct BIRC5^+^ cluster, separate from both the myeloid and lymphoid clusters (Figure 2B). However, distinguishing cells derived from either WT- or ΔTOX-infected mice revealed that it was only in mice infected with the WT strain of *S. aureus* that this BIRC5^+^ population was elicited (Figure 2C,D). There were other minor differences, with slightly different proportions of the monocyte/macrophage clusters, and a greater percentage of cells characterized as NK cells in the ΔTOX-infected mice (Figure 2D). The BIRC5 cluster can be identified as myeloid cells based on expression of Ms4a3 (Figure 2E). Some NKT cells that express Cd3d also express Birc5, but they fall clearly into the lymphoid clusters (Figure 2E). Within the BIRC5^+^ cluster, differential analysis of gene expression between cells from WT and ΔTOX-infected mice showed that the few cells in this cluster from ΔTOX-infected mice express lower levels of inflammation related genes such as Ccl4, Mif and Ctsg, as well as Ms4a3 (Figure 2F). These data suggest that the elicitation of a population of BIRC5^+^ myeloid cells in the liver during *S. aureus* infection is dependent on the production of leukotoxins by the bacteria.

### BIRC5^+^ immune cells share a common transcriptional program across a range of inflammatory states

To determine if BIRC5^+^ myeloid cells derived from CX3CR1^+^ precursors are present across a variety of inflammatory states, we integrated scRNA-seq datasets from several fate-mapping experiments that we have performed. The combined dataset consisted of data from CD45^+^ tdTomato^+^ cells isolated from the liver of *S. mansoni* and *S. aureus* infected mice as described above (Figure 1 and Figure 2), a subset of liver CD11b^+^ tdTomato^+^ cells isolated during *S. mansoni* infection, and a previously published dataset in which BIRC5 monocytes were initially identified in the aorta from a model of atherosclerosis (3). After integration and dimension reduction, we visualized the cells according to their transcriptional profile and the dataset of origin (Figure 3A). Many of the clusters corresponded to the original datasets, for example cluster 1 consisted mostly of cells from mice infected with *S. aureus*, although other datasets are represented in this cluster as well (Figure 3B). Cells derived from *S. mansoni* infected mice appeared to be the most transcriptionally distinct based on the clustering. The expression of BIRC5 was concentrated in clusters 6 and 9, each of which consisted of cells drawn from multiple datasets, suggesting that the BIRC5^+^ population has a common transcriptional profile across multiple inflammatory conditions (Figure 3C-E). These same clusters had higher expression of Ki67 and STMN1, indicating a shared capacity for proliferation. Data from this experiment can be explored at https://ruggleslab.shinyapps.io/ProliferatingMonocytes/.

**Figure 3.**
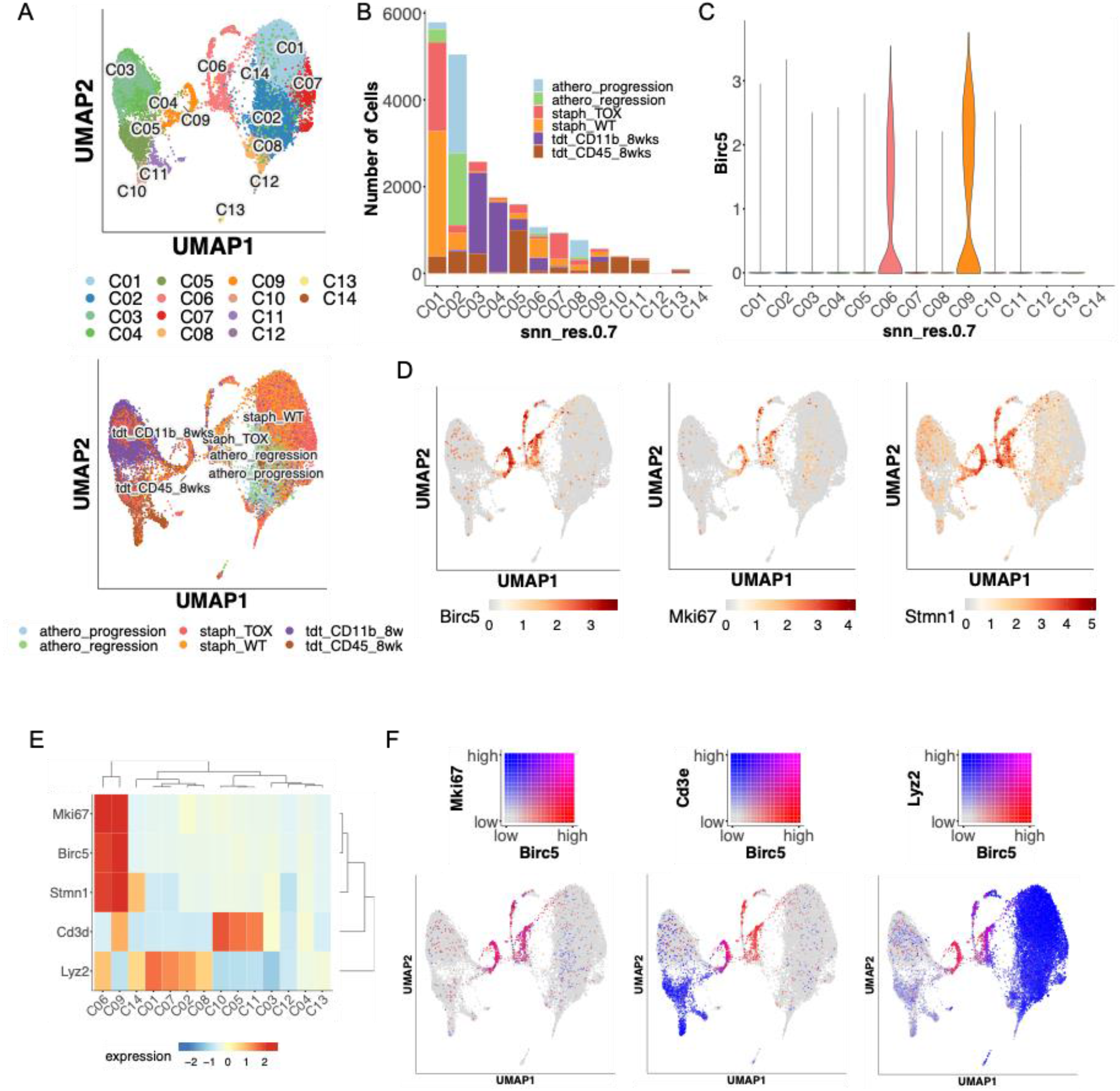
Combined UMAP clustering of single-cell RNA-sequencing datasets reveals common transcriptional profile of BIRC5^+^ immune cells across multiple inflammatory states. **(A)** UMAP clustering of CX3CR1^+^-derived cells from multiple datasets based on transcriptional profile and experimental origin. **(B)** Number of cells present in each defined cluster, demonstrating common transcriptional profiles across multiple inflammatory settings. **(C)** Violin plot of *BIRC5* expression. (**D)** Gene expression levels of *BIRC5* coinciding with expression of the proliferation markers *MKI67* and *STMN1*. **(E)** Heatmap of relevant gene expression across all clusters. **(F)** Coexpression of *BIRC5* with *MKI67, CD3e*, and *LYZ2*, showing *BIRC5* expression in both a lymphoid and a myeloid population. Figures generated using SHINY app (ruggleslab.shinyapps.io)

Utilizing the Shiny application, we generated gene pair heat plots to visualize overlaps in expression. Comparing BIRC5 to Ki67 expression, we see near complete overlap, indicating the BIRC5^+^ population as the major proliferating population across all datasets (Figure 3F). Similar to what was observed in the individual analyses, we found two clusters of BIRC5^+^ cells, one lymphoid and the other myeloid, as indicated by the overlap between the genes CD3e and Lyz2, respectively, and BIRC5 expression (Figure 3F). Cluster 6 corresponded to the BIRC5^+^ lymphoid cluster while cluster 9 corresponded the BIRC5^+^ myeloid cluster. BIRC5^+^ myeloid cluster is also marked by expression of Ms4a3 and other myeloid markers. Overall, combined analysis revealed BIRC5^+^ immune cells share a transcriptional profile across multiple inflammatory states, with a clear distinction between BIRC5^+^ myeloid cells and BIRC5^+^ lymphocytes.

### Conditional deletion of BIRC5 in CX3CR1^+^ cells improve survival during *Staphylococcus aureus* infection

To further assess the role of BIRC5^+^ immune cells *in vivo*, we crossed mice in which the *BIRC5* gene was flanked by two loxP sequences to the CX3CR1^CreERT2-IRES-eYFP^ mice, so that when tamoxifen is administered, the *BIRC5* gene is conditionally deleted. Progeny were then crossed to generate mice homozygous for both alleles. We infected double homozygous mice, along with mice homozygous for either the CX3CR1^CreERT2-IRES-eYFP^, or the BIRC5-Flox, alone, with 1×10^7^ CFU of *S. aureus*, a dose known to cause significant mortality in mice. To induce deletion, mice were fed chow containing tamoxifen, starting 2 days prior to infection, up until the end of monitoring at day 10 post-infection. Mice were then monitored for weight loss and survival. We observed significantly less mortality in the double homozygous group as compared to other groups, suggesting that deletion of BIRC5 in CX3CR1^+^ cells is protective against lethality (Figure 4A). Approximately 50% of mice in the double homozygous group survived to day 10 whereas only 5-15% of mice in the control group survived until day 10. Furthermore, mice in the control group started to die sooner during the course of infection, mostly between days 2 and 4. Observing this difference in survival, we assed bacterial burden in both CX3CR1^+/CreERT2-IRES-eYFP^ BIRC5-Flox, and BIRC5-Flox alone mice after infection with 1×10^7^ CFU of *S. aureus*. All mice were placed on tamoxifen diet 2 days prior to infection, and at day 2 post-infection, liver, spleen, heart, kidney, and lung were harvested to quantify bacterial burdens. There was a lower bacterial burden in the livers and spleens of mice in which BIRC5 was conditionally deleted in CX3CR1-expressing cells, with no significant difference in the kidney, heart, and lung (Figure 4B, data not shown). These results show that the deletion of BIRC5 in CX3CR1^+^ cells protect mice from acute lethality from *S. aureus* infection, with reduced bacterial burden in the liver and spleen. As this difference occurs within the first 3 days of infection, the results implicate CX3CR1^+^ myeloid cells as playing a pathogenic role in increasing bacterial burden in the liver during acute *S.aureus* infection.

**Figure 4.**
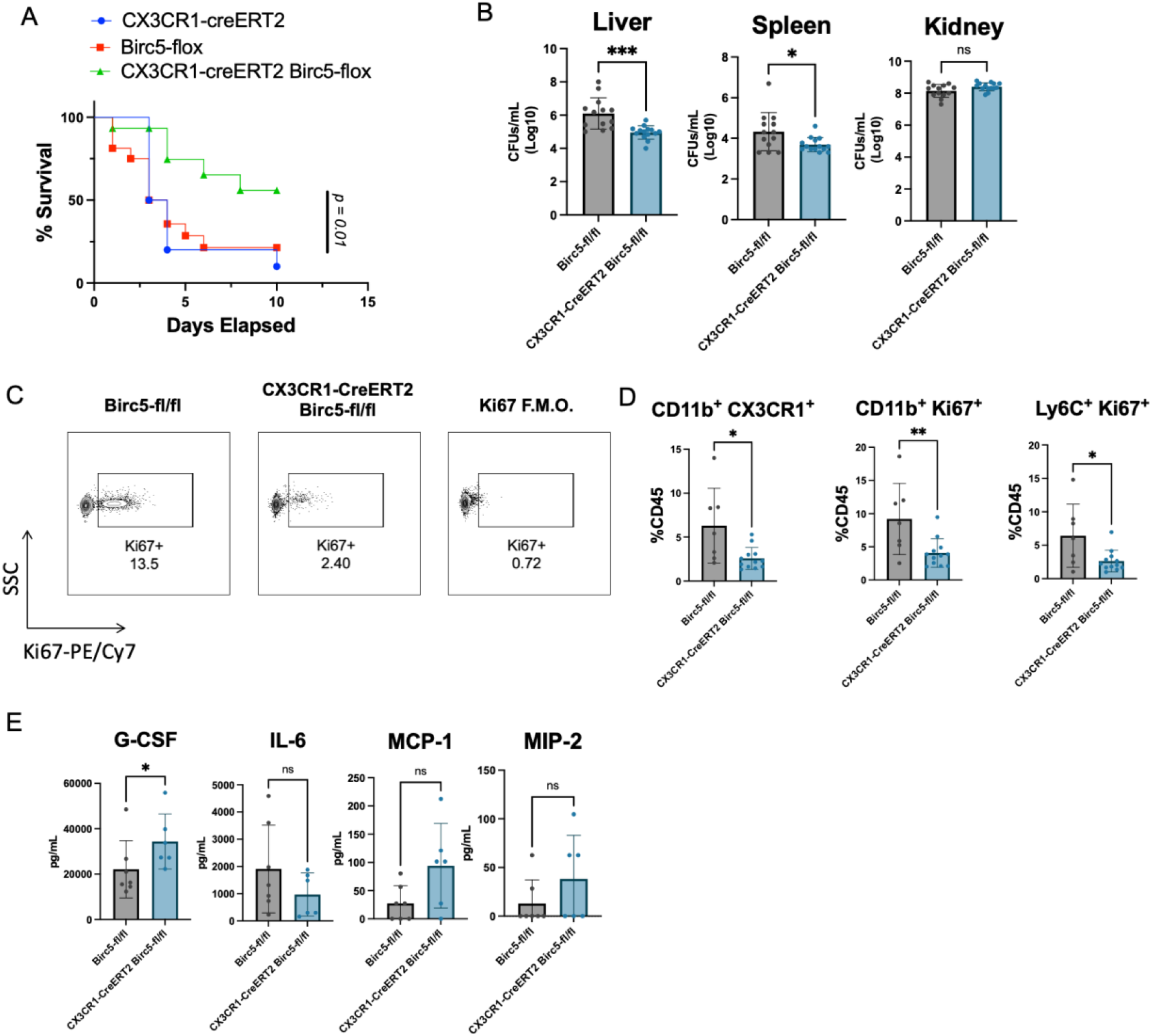
Conditional deletion of BIRC5 in CX3CR1-expressing cells is protective during infection with *S. aureus*. **(A)** Survival curve of mice infected with 1×10^7^ CFU of *Staphylococcus aureus* (n = 13/14 mice per genotype). **(B**) Bacterial burden across tissues at day 2 post-infection, with values log transformed for standardization (n = 12-14 mice per genotype). **(C)** Representative flow cytometry plots of Ki67 expression in CD11b^+^ leukocytes in the liver at day 2 post-infection, gated on Live cells, CD45^+^,CD11b^+^. **(D)** Percentage of monocyte populations as part of total CD45^+^ at day 2 post-infection in the liver (showing combination of 2 different experiments, n = 7-12 mice per genotype). **(E**) Concentration of cytokines/chemokines per milliliter of blood was quantified by the magnetic bead assay for blood infected from either Birc5-Flox of CX3CR1-creERT2 Birc5-Flox after sera was harvested at day 2 post-infection (n = *6⊓* per genotype). Error bars represent SEM. Student’s *t* tests were performed to determine significance. *P<0.05; **P<0.01; ***P<0.001; ****P<0.0001.

To assess the immune response in these mice, flow cytometry was performed on cells isolated from the liver. After gating on CD45^+^ CD11b^+^ cells, we observed a lower incidence of Ki67 expression in mice in which BIRC5 was conditionally deleted, suggesting that BIRC5 is needed for proliferation of the myeloid cells in the liver (Figure 4C). Quantifying cells as a percentage of total CD45 cells showed there to be a significant reduction in proliferating monocytes upon deletion of BIRC5 (Figure 4D). We next determined if deletion of BIRC5 from CX3CR1^+^ cells resulted in altered cytokine and chemokine profiles at a systemic level. We noticed very little difference between the two genotypes, with the most notable difference being a modest increase in the levels of G-CSF in mice in which BIRC5 was conditionally deleted (Figure 4E). Although not statistically significant, there was a trend toward higher levels of MCP-1 (CCL2) and MIP-2 in the conditional knockout mice (Figure 4E). Inflammatory cytokines showed little difference, although there was trend toward lowers levels of IL-6 in the conditional knockout (Figure 4E). Overall, these data show that the deletion of BIRC5 in CX3CR1^+^ cells result in reduced proliferation within the monocyte population as well as lower numbers of total monocytes in the liver, but with minor effects on other immune cell populations and inflammatory parameters.

## Discussion

In this study, we determined that a population of BIRC5^+^ cells derived from CX3CR1^+^ precursors is a general feature of both acute and chronic infections in the liver, during both bacterial and helminth infections. While there are both BIRC5^+^ lymphocytes and BIRC5^+^ myeloid cells, the deletion of BIRC5 from CX3CR1 expressing cells results in improved survival during acute *S. aureus* infection. The 3-day time scale implicates BIRC5 in the myeloid cell compartment as playing a pathogenic role during *S. aureus* infection, by promoting bacterial burden in the livers of infected mice. In the absence of toxins, *S. aureus* infection does not elicit these BIRC5^+^ cells, implicating a toxin dependent pathogenic role for BIRC5 in myeloid cells during *S. aureus* infection.

As part of this study, we have further defined the subsets of immune cells that express the chemokine receptor CX3CR1 using a combination of fate-mapping mouse models and single-cell RNA-sequencing. We found this receptor to be expressed on multiple immune cell types, including members of both the myeloid and lymphoid lineages, demonstrating a greater level of diversity in expression than previously realized. For our infection studies we focused on a single tissue – the liver – but compared disparate infections to determine how immune cells respond to diseases with categorically different pathogen biology and etiologies. During infection with the trematode parasite *S. mansoni*, the liver is the main site of pathology, and the immune response characterized by the infiltration of monocytes to form a granuloma structure around the parasite eggs (4–6). During infection with the Gram-positive bacteria *S. aureus*, the liver is also one of the primary tissues affected, as the bacteria is sequestered in the liver within hours of an intravascular infection (26,27). By conducting our analysis on both a ‘Type 2’ and ‘Type 1’ infection respectively and combining this with our analysis of a sterile inflammatory setting – atherosclerosis, we were able to identify transcriptional signatures, both common and distinct, of CX3CR1-expressing immune cells responding to inflammation. Most notably was our identification of a broadly occurring BIRC5^+^ population comprising both lymphoid and myeloid cells exhibiting stem cell – like properties. Numerous studies have demonstrated a role for BIRC5^+^ T cells in infection and autoimmunity, however our study is unique in demonstrating the occurrence of a BIRC5^+^ myeloid population during multiple inflammatory settings as well as showing this population to be pathogenic during *S. aureus* infection.

Due to its role in cell cycle progression, BIRC5 has been hypothesized to be an important biomarker of many cancers. Indeed, multiple studies have shown a positive correlation between expression levels of BIRC5 in different tumors and a poorer prognostic outcome (15,16,29–31). Furthermore, while not true for all cancers, many tumors displayed a positive correlation between BIRC5 expression and immune cell infiltration, suggesting a potential immunogenic role for BIRC5 (15,31). These data suggest BIRC5 could be a therapeutic target in certain tumor settings although the efficacy of such a therapy remains to be tested. However, our own data suggests that many immune cells themselves would be targeted by such a therapy, and so it is crucial to take this into consideration assessing the potential for targeting BIRC5.

One limitation of this study is that it is unclear why conditional deletion of BIRC5 in CX3CR1^+^ cells is protective during *S. aureus* infection. It is known that tissue resident macrophages in the liver (Kupffer cells) will rapidly sequester the bacteria upon intravascular infection (26,27). During the first 24 hours of infection, many of these cells will manage to kill the bacterium, however a significant portion will instead be overwhelmed by bacterial replication, subsequently lysed, and the bacteria allowed to disseminate. It is only after this initial phase of infection in the liver that the bacteria become systemic, spreading to other tissue and the blood. Recent work has shown that after the liver phase of the infection, the bacteria spreads to the peritoneal cavity where it is taken up by a population of GATA6^+^ macrophages before spreading widely (28). However, the same study showed multiple cell types take up the bacteria, including monocytes both in the liver and peritoneal cavity. In the context of our own study, it is possible that by deleting BIRC5 in CX3CR1^+^ cells we are reducing the cellular hosts for the bacteria, which prevents uncontrolled growth and in turn results in better survival. It has been shown that monocytes are able to engraft in the liver to replace depleted Kupffer cells, so deletion of BIRC5 in monocytes could prevent the replenishment of these macrophages and starve the bacteria of cellular hosts after the initial wave of lysis and dissemination (32).

Furthermore, the reduction in monocytes upon conditional deletion could be what is causing increased systemic levels of G-CSF and CCL2 as there is an open niche for recruitment of new myeloid cells. Given the wide range of tissues affected by infection and the many different myeloid cells present in said tissues, it is possible that conditional deletion has a stronger effect in some tissues over others. We observed lowered bacterial burden in the liver and spleen but not in the kidney, lung, or heart after 48 hours, suggesting that we are not preventing spread to other tissues but arresting the growth of bacteria in the liver and spleen specifically. Remarkably, this was enough to improve survival.

Another possibility that we did not fully explore in this study is whether CX3CR1^+^ BIRC5^+^ immune cells are regulatory in nature. It is known that *S. aureus* infection can actually suppress the host immune system in ways that are still being determined but include toxin-mediated killing of leukocytes (9,10,12). One interesting observation from our study was that infection with the ΔTOX strain of bacteria did not elicit the BIRC5^+^ population in the liver. Since the absence of this population in our genetic models improved survival, it could be mimicking an infection seen with a ΔTOX strain of the bacteria where lethality is reduced (11,39). This phenotype could be mediated by toxin killing of immune cells or simply bacterial toxins inducing a dampened immune response, unable to strongly respond to infection (12). With the current lack of vaccines to protect against *S. aureus* infections as well as the development of antimicrobial resistant strains, it is imperative to pursue alternate treatment options. One such approach that has gained traction against infectious disease has been host directed therapies, whereby the host cellular processes needed by the pathogen lifecycle are disrupted (33). Other studies have shown this approach to be effective by, for example, disrupting the apoptosis pathway (34,35). Our study has revealed a new possible target for host directed therapy in BIRC5, as conditional deletion of this gene in immune cells significantly improved survival. Further understanding of the role of BIRC5^+^ immune cells during infection with *S. aureus* is crucial for determining the possibility of targeting this protein therapeutically.

## Author contributions

Conceptualization: DGL, JCD, JL, VJT, PL. Methodology: DGL, JCD, KAL, JL, VJT, PL. Investigation: DGL, JCD, KAL, EEZ, JL, PL. Data curation and analysis: DGL, JCD, KAL, NH, ZC, JL, KVR, PL. Writing – original draft: DGL, PL. Writing – review and editing: DGL, VJT, PL. Visualization: DGL, JCD, NH, ZC, JL, KVR, PL. Supervision: KMK, VJT, PL. Funding Acquisition: KMK, VJT, PL.

## Acknowledgements

We acknowledge help from the NYU Langone’s Genome Technology Center for performing all the RNA sequencing. Cell sorting/flow cytometry technologies were provided by NYU Langone’s Cytometry and Cell Sorting Laboratory. E.E.Z. was supported in part by National Institute of Allergy and Infectious Diseases (NIAID)–Supported Institutional Research Training Grant on Infectious Disease & Basic Microbiological Mechanisms T32 AI007180. J.C.D. was supported in part by NIAID-Supported Institutional Research Training Grant in Immunology and Inflammation T32 AI100853. K.A.L. was supported by Cystic Fibrosis Postdoctoral Research Fellowship Award LACEY19FO. J.L is supported by Young Scholar Fellowship-Einstein Grant and Yushan Scholar Program, from the National Science and Technology Council and the Ministry of Education, Taiwan. This research was also supported in part by the Intramural Research Program of the NIAID, NIH (P.L.). Research in the laboratory of V.J.T. is supported by NIH, NIAID Awards AI099394, AI105129, AI137336, AI133977 and by a Burroughs Wellcome Fund Investigator in the Pathogenesis of Infectious Diseases award. The NYU Langone Health Genome Technology Center and the Cytometry and Cell Sorting Laboratory are shared resources partially supported by Laura and Isaac Perlmutter Cancer Center Support Grant P30CA016087 from National Institutes of Health The National Cancer Institute (NIH-NCI).

